# Identification of targetable epitope surfaces from the high resolution structure of the superantigen *Staphylococcal* Enterotoxin L

**DOI:** 10.64898/2026.03.13.711576

**Authors:** Ella Klamm, Caterina Curato, Katherina Siewert, Stephen F. Marino

**Affiliations:** Department of Biological Safety, Department of Chemical and Product Safety of the German Federal Institute for Risk Assessment, Berlin, Germany; Dermatotoxicology Study Centre, Department of Chemical and Product Safety of the German Federal Institute for Risk Assessment, Berlin, Germany

**Author notes:** For Correspondence: Stephen F. Marino.

**Keywords:** bacterial toxin, superantigen, X-ray crystallography, *Staphylococcus aureus* (*S. aureus*), T cell receptor (TCR)

## Abstract

Emetic exotoxins secreted by *Staphylococcus* species (*Staphylococcal* enterotoxins, SEs) are a major cause of food poisoning cases worldwide and many are additionally classified as superantigens – able to potently activate T cells in an antigen independent manner. Fewer than half of the gene products of the known SE genes have been extensively characterized. The gene for Staphylococcal enterotoxin L (SEL) occurs in both foodborne and clinical isolates but no detailed structural characterization has yet been available. We report here the crystal structure of SEL and confirm its function as a superantigen via direct T cell activation assays. By comparison of the SEL sequence with that of its four closest homologues (SEI, SEK, SEM and SEQ), we have identified binding epitopes unique for SEL and mapped these regions onto the structure. These data provide the first high resolution view of SEL and the basis for the development of diagnostic procedures for its specific detection.

## Introduction

Bacteria in the genus *Staphylococcus* are common colonizers of both livestock and humans and antibiotic resistant strains such as MRSA (methicillin resistant *S. aureus*) are frequent and dangerous causes of nosocomial infections worldwide. Among other virulence factors, *Staphylococcal* sp., primarily *S. aureus*, harbor an array of genes for secreted toxins termed SEs (*Staphylococcal* enterotoxins) whose known differentiable sequences currently number about 30 [1]. Their gene products are relatively small (27 – 30 kD) compactly folded proteins that, at least for those that have been extensively studied, show high resistance to proteases, extremes of pH and heat, enabling them to easily persist under conditions that render the bacteria inactive. Most characterized SEs (named according to letters of the English alphabet in the order they were discovered, SEA, SEB, etc.) induce severe emesis upon ingestion. SEs are thereby among the leading causes of strong evidence foodborne outbreaks (FBOs) annually in the EU and their incidence is increasing [2]; they are additionally expected to play a substantial role in the nearly one-third of food poisoning cases each year for which no cause can be determined. Many emetic SEs are also classified as superantigens (SAgs) because they are able to simultaneously bind MHC II (major histocompatibility complex II) molecules and TCRs (T cell receptors) resulting in the activation of T cells independent of MHC II bound peptide [3]. This subversion of the normally strictly regulated T cell activation process is so efficient that the presence of superantigenic SEs in the system can provoke a several thousand-fold higher activation of the total T cell population than experienced during a normal immune response – the resulting massive release of cytokines from these activated T cells, a so-called cytokine storm, can lead to the potentially fatal Toxic Shock Syndrome.

Crystal structures have demonstrated that these toxins share similar topologies despite, in some cases, substantial sequence variability [3]. There is nevertheless significant sequence similarity shared among some groups of SEs, a fact that severely hampers the development of specific detection methods for their rapid identification in food samples and in the clinic. Considering the increasing incidence of MRSA cases [4] and the continuing discovery of new SE genes, the need for specific detection procedures – and for specific therapeutic options to block their superantigenic activities – is steadily growing.

SEs have been classified into groups (I – V) based on their sequence similarities and corresponding biochemical properties. Despite the divergence that exists between these different groups, the overall SE sequence similarities make their specific detection in foodstuffs or clinically derived material extremely difficult. The standard and most cost effective method employed, that of antibody-based ELISAs, depends for its specificity on the recognition of unique epitopes on the surfaces of the antigens. However, in the absence of three-dimensional structures of the toxins of interest, the identification of unique epitopes for the targeted development of specific recognition units, whether IgGs or designed binding modules, remains a formidable challenge.

The gene for SEL, initially identified in an *S. aureus* isolate from a bovine mastitis case [5], has been subsequently identified in many isolates from both clinical and foodborne sources but there has been no detailed structural characterization of the protein. Previous *in vitro* interaction studies monitoring cytokine release from toxin treated mouse splenocytes [6] as well as detection of TCR-β chains on toxin activated T-cells [7] indicated that SEL can function as a superantigen but animal studies demonstrated only minimal emetic activity.

We therefore undertook to produce SEL recombinantly for further characterization of this toxin and, in order to verify its wildtype function, tested our purified material *in vitro* via human T cell activation experiments. We report here the three-dimensional structure of SEL and confirm its classification as a superantigen. Further, we identify regions on the molecule’s surface that can be targeted for the development of specific binders able to distinguish SEL from other Group V SEs.

## Results

### Structure and details

We solved the x ray structure of SEL to 1.3 Å resolution (Fig S1, Table S1) and the visible density covers the entire wildtype amino acid sequence except for the N-terminal asparagine residue (residues 2 – 216). The wildtype secretion signal sequence was substituted with N-terminal His_6_ and FLAG tags for purification but no density is observed for any of these residues. The SEL structure shows the typical SE topology, comprising an N-terminal β-barrel and a C-terminal β-grasp domain. According to the structure based SE classification of Spaulding, *et al*. [8], SEL had been assigned to the Group V SEs. The core trace of SEL mirrors that shared by the only other Group V SEs for which structures are available, SEK (in complex with TCRβ 5.1 (PDB ID: 2NTS) [9]) and SEI (in complex with a peptide-loaded MHC II (2G9H) [10]) and aligns with SEK with a mainchain rmsd of 2.1 Å (residues 2 – 216), and with SEI with a mainchain atom rmsd of 3.2 Å (residues 4 - 216)(Fig 1). (A structure of SEK alone is also available (3EA6) and aligns with an all atom rmsd of 1.2 Å (residues 1 – 217) to the structure of the TCRβ-complexed toxin; when not otherwise stated, all comparisons with SEL reported here are with the two complex structures).

**Fig. 1:**
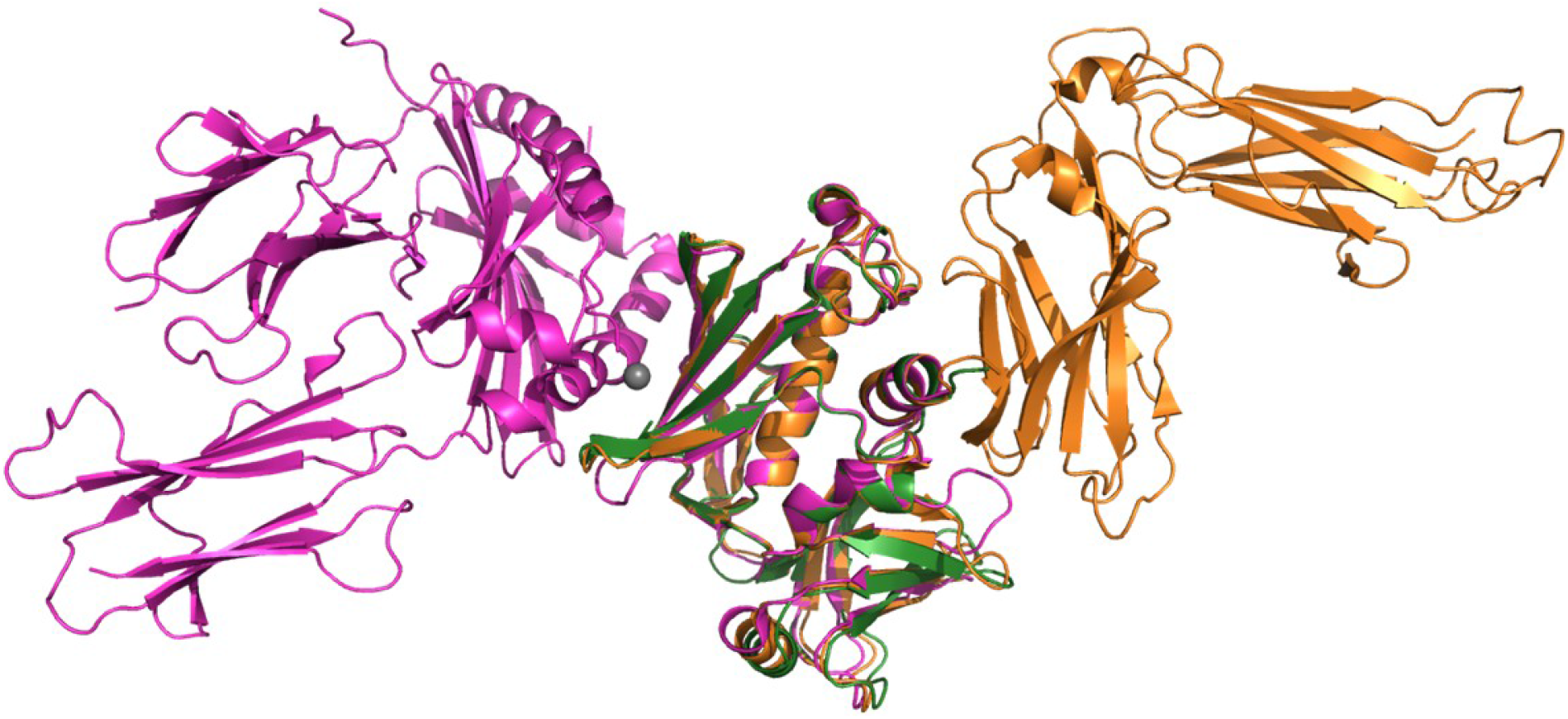
Structural alignment of SEL with SEI (2G9H, in complex with MHC II) and SEK (2NTS, in complex with TCRβ5.1). Alignment of the mainchain atoms of SEL (green) with those of SEI (magenta) and SEK (orange) depicting the minimum T cell activation complex. The MHC II molecule from the SEI complex is shown on the left side of the aligned toxins in magenta while TCRβ5.1 from the complex structure with SEK is shown in orange on the right side.

The largest topological deviations between the structures are confined to three loop regions; at two, SEL and SEK correspond well but differ from SEI (residues 68 – 76 and residues 170 – 176) while the third shows an SEL:SEI correspondence deviating from SEK (residues 143 – 149) (Fig 2). The loop comprising residues N70 – S75 in SEI is markedly displaced from the position of the analogous loop in both SEK and SEL, due in part to the presence of Pro72 in the center (a Gly residue occurs at this position in SEL and a Gln in SEK). Residues from this loop do not contact the TCR in the SEK complex structure, but the substantial outward displacement seen in the SEI structure could indicate its participation in TCR binding in that case.

**Fig 2:**
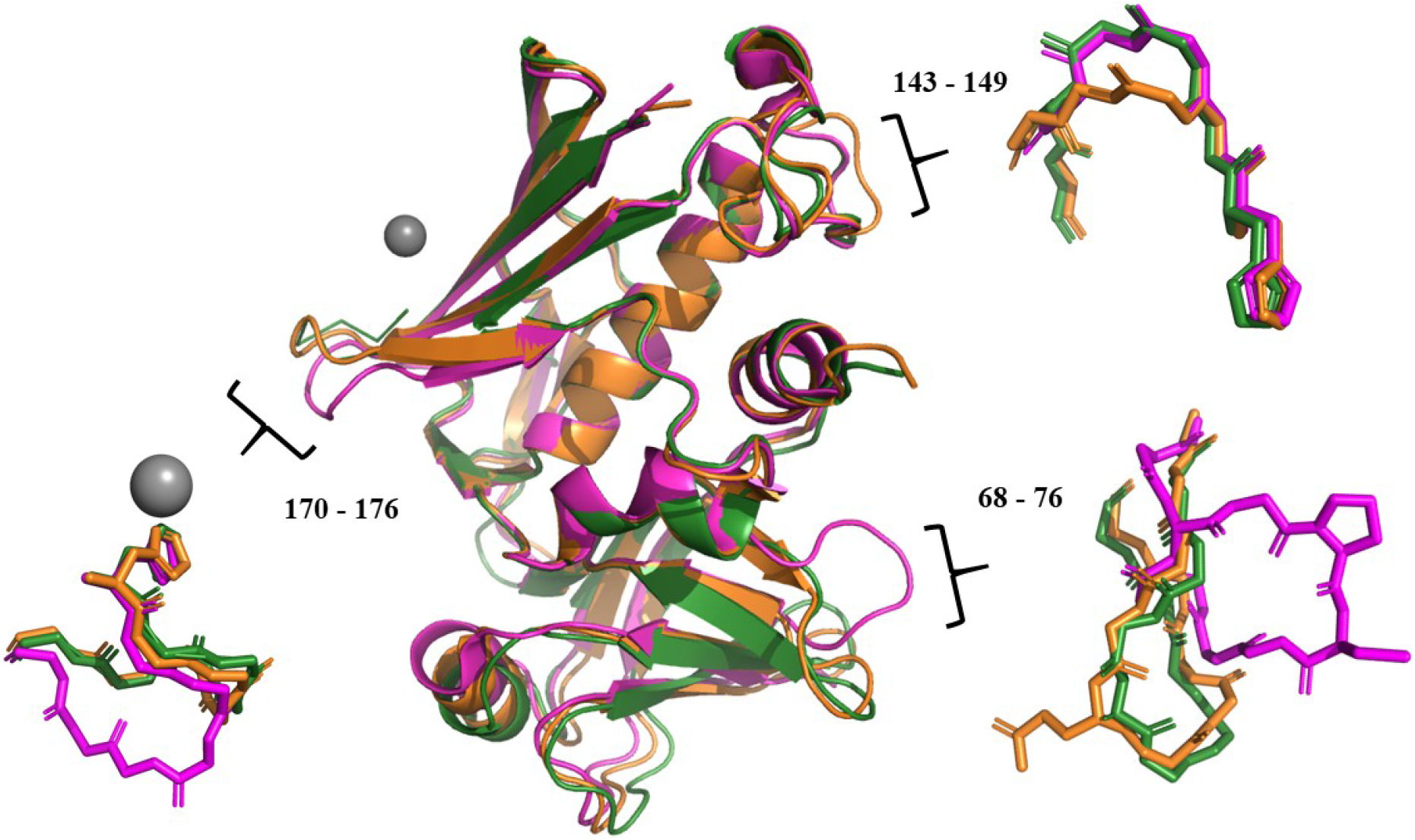
Structural deviations between SEL and its close homologues SEI and SEK. Mainchain atom alignment of the three toxins (SEL in green, SEI in magenta and SEK in orange) depicted as ribbons indicating the three divergent loop regions. The isolated loops are shown in stick representation. SEI deviates substantially in loop regions 68 – 76 and 170 – 176 from the well aligned traces of SEK and SEL, while the loop comprising residues 143 – 149 in SEK shows a slight deviation from the nearly identical orientation of SEI and SEL in this region.

The SEL α3β8 loop (comprising residues 141 - 160), a sequence insertion characteristic of Group V SEs and participating in interactions with TCRβ chains, largely follows the trace of the corresponding loop visible in the available structures of SEK with the positions of the flanking sidechains H142 and Y158 being essentially identical. The alignment of this region shows no difference in the positions of the loop in the two published SEK structures [9] (Fig 3, upper left panel). The stretch between residues 143 and 149 does however show two differences between SEL and SEK, the first being the sidechain of the boundary Y159 in SEL whose orientation deviates substantially from that of the corresponding Y158 in SEK (Fig 3, right panel). As Y159 in SEL participates directly in a crystal contact with a neighboring SEL molecule, the observed position of this side chain is unlikely to have any physiological significance. The second deviation concerns the region encompassing residues N144 and K148 in the center of the α3β8 loop (Fig 3 lower left and right panels). The position of this peptide is shifted by nearly 3 Å from that in SEK (measured between the alpha carbon atoms of Thr145 in SEK and Thr146 in SEL), in part due to the flipping of the peptide bonds between residues Asn144 and Gly145 and Thr145 and Lys146, but adopts a position nearly identical to that of the corresponding peptide in SEI. It Is possible that this difference is related to TCRβ chain preference.

**Fig 3:**
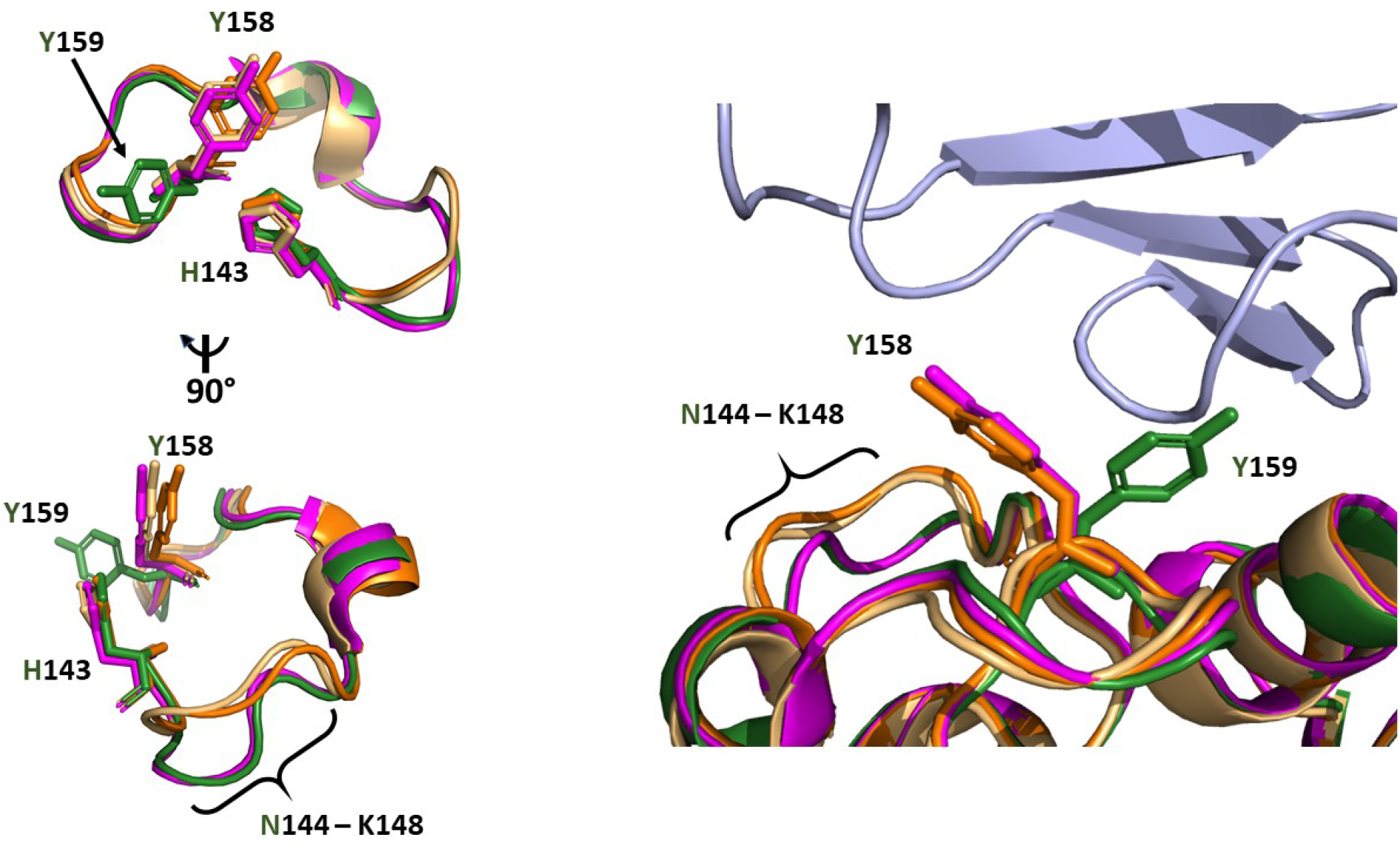
Structural alignment of the α3β8 loop. Structural alignment of the SEL (green), SEI (magenta) and SEK (orange) structures - the structure of SEK alone (3EA6) is also included here in light orange for comparison – showing the α3β8 loop region. The loop boundary residues are explicitly labelled in the two views in the left panel as well as the side chains of Y158 and Y159 in the right panel. The binding interface region of TCRβ5.1 from the SEK complex structure (2NTS) is shown in blue in the right panel. The altered conformation of the N144 – K148 loop in SEK compared to that of SEI and SEL is marked in both panels.

The loop comprising residues 170 – 176 adopts a conformation nearly identical to that seen in SEK but the corresponding loop in SEI is displaced away from the MHC II binding site, with none of the residues being involved in the complex interaction. Considering the relative position of this loop in SEL and SEK, it may form part of the MHC II interface with these toxins.

One further segment of the SEL structure that deviates from that observed in both SEI und SEK comprises residues 84 - 93 and 114 – 117 in the latter two toxins at the base of the C-terminal β-grasp domain. These stretches form a two stranded β-sheet in SEI and SEK while the equivalent strands in SEL, 88 – 94 and 115 – 118, do not reflect the canonical pairing parameters and are modelled as loops (Fig 4A). In none of the three structures do these residues participate in any binding interactions but they are adjacent to the interface with the MHC II molecule in the SEI complex structure, where Lys94 of SEI forms a salt bridge with Glu69 of the MHC II. The proximity of this region to the MHC II interface suggests the possibility of differences in this interaction between SEI and SEL/SEK.

**Fig 4:**
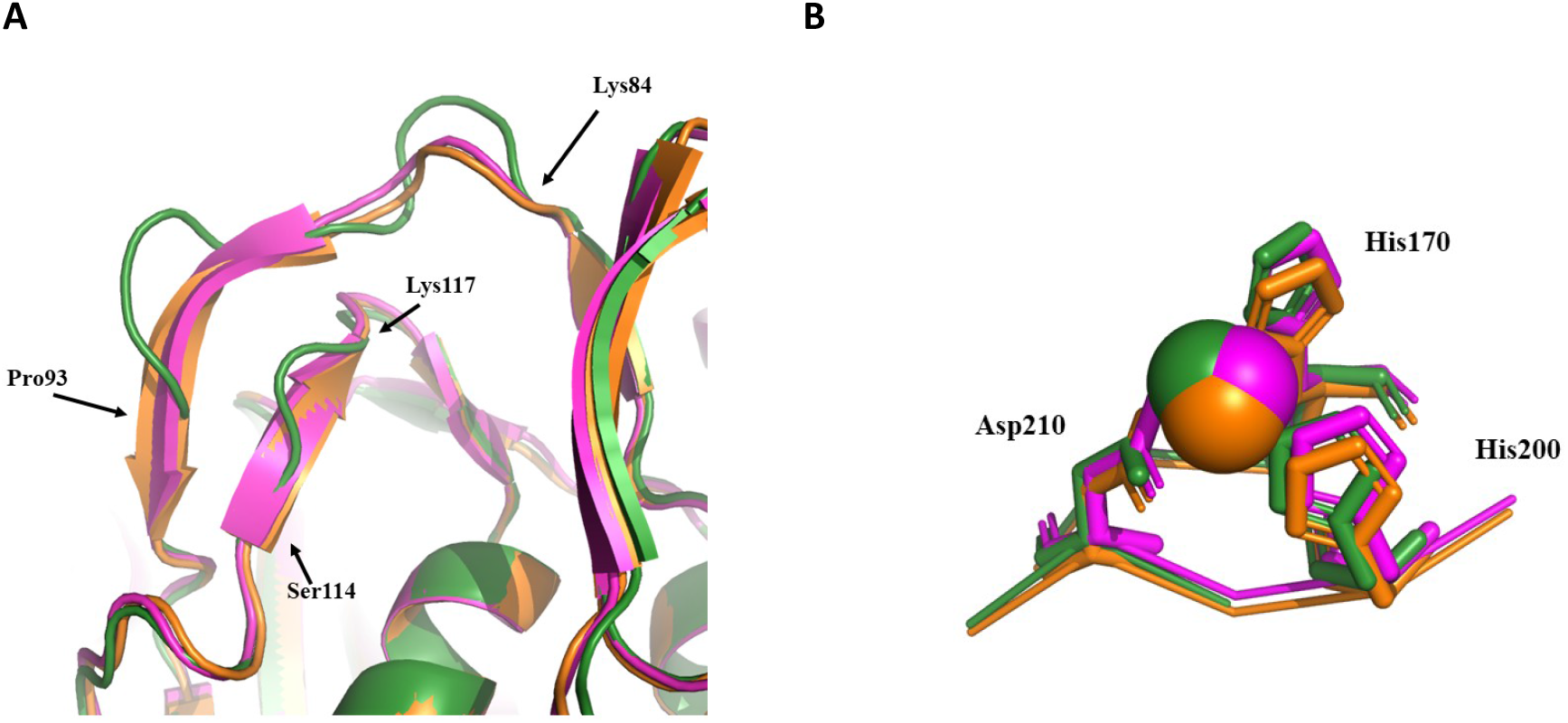
Structural alignment comparing the two-stranded β-sheet of the C-terminal domain. **A**. The base of the β-grasp domain in SEL (green) SEI (magenta) and SEK (orange) is depicted. SEI and SEK both display a nearly identically positioned two-stranded β-sheet leading into the C-terminal domain, the same region in SEL does not. This difference may indicate an altered MHC II binding interface for SEL. **B**. The Zn^++^ binding site of SEL. Mainchain atom alignment of SEL (green), SEI (magenta) and SEK (orange) depicting the Zn^++^ binding site in each; the Zn^++^ atom is colored in each case according to its respective structure.

A primary feature of the Group V SEs is the presence of a Zn^++^ binding site located on the MHC II binding face of the protein. A single Zn^++^ ion is visible in the SEL structure, coordinated by residues His170, His208 and Asp210, whose positions show excellent agreement with the analogous binding sites in the available structures of SEK and SEI (Fig 4B). While an MHC II molecule provides a His residue as the fourth protein ligand for the bound Zn^++^ in the SEI complex structure, in SEL this His residue (His 143) is provided by a neighboring monomer in the crystal.

### Superantigen-mediated T cell activation by SEL

We employed a CD154-based activation-induced marker (AIM) assay to assess T cell reactivity to SEL [11, 12]. Human PBMCs were incubated with a 10-fold concentration series of SEL for 5 hours (10 ^-9^ – 10 μg/mL). SEB, a well-studied superantigen, served as the positive control and was analysed in parallel. T cell reactivity was assessed by flow cytometry, monitoring the induced upregulation of activation markers including CD154 (CD40L), CD69 and CD137. Surface staining revealed a prominent upregulation of CD154 on CD4+ memory T cells for both SEL and SEB (Fig 5; for the full gating strategy, see Fig S2). At the highest SAg concentrations, up to ∼30% CD154+CD4+ memory T cells were observed. Example dot plots and summarized data illustrate that the frequencies of activated CD154+CD4+ memory T cells were markedly higher at lower concentrations of SEL than for SEB (Fig 5). For instance, at 1 pg/mL, 3.04% vs. 0.21% of activated CD4+ memory T cells were observed in a representative experiment for SEL and SEB, respectively (Fig 5A). Similar values were obtained in four independent experiments (Fig 5B). Expression of additional T cell activation markers CD69 and CD137 (4-1BB) confirmed this trend (Fig S3 A-B). Compared to SEB, SEL showed no interaction with TCR β-chain vβ17 and neither superantigen showed interaction with vβ 13 (Fig S2 and not shown). Superantigen-mediated activation was also observed for CD4+ naïve T cells and CD8+ memory T cells, utilizing CD154 and CD137 as activation markers, respectively (Fig S3 C-D). The CD137+CD8+ memory T cell population showed no difference for SEL and SEB, which may be linked to both a higher background expression and possible share of non-specific induction of this marker [13].

**Fig 5:**
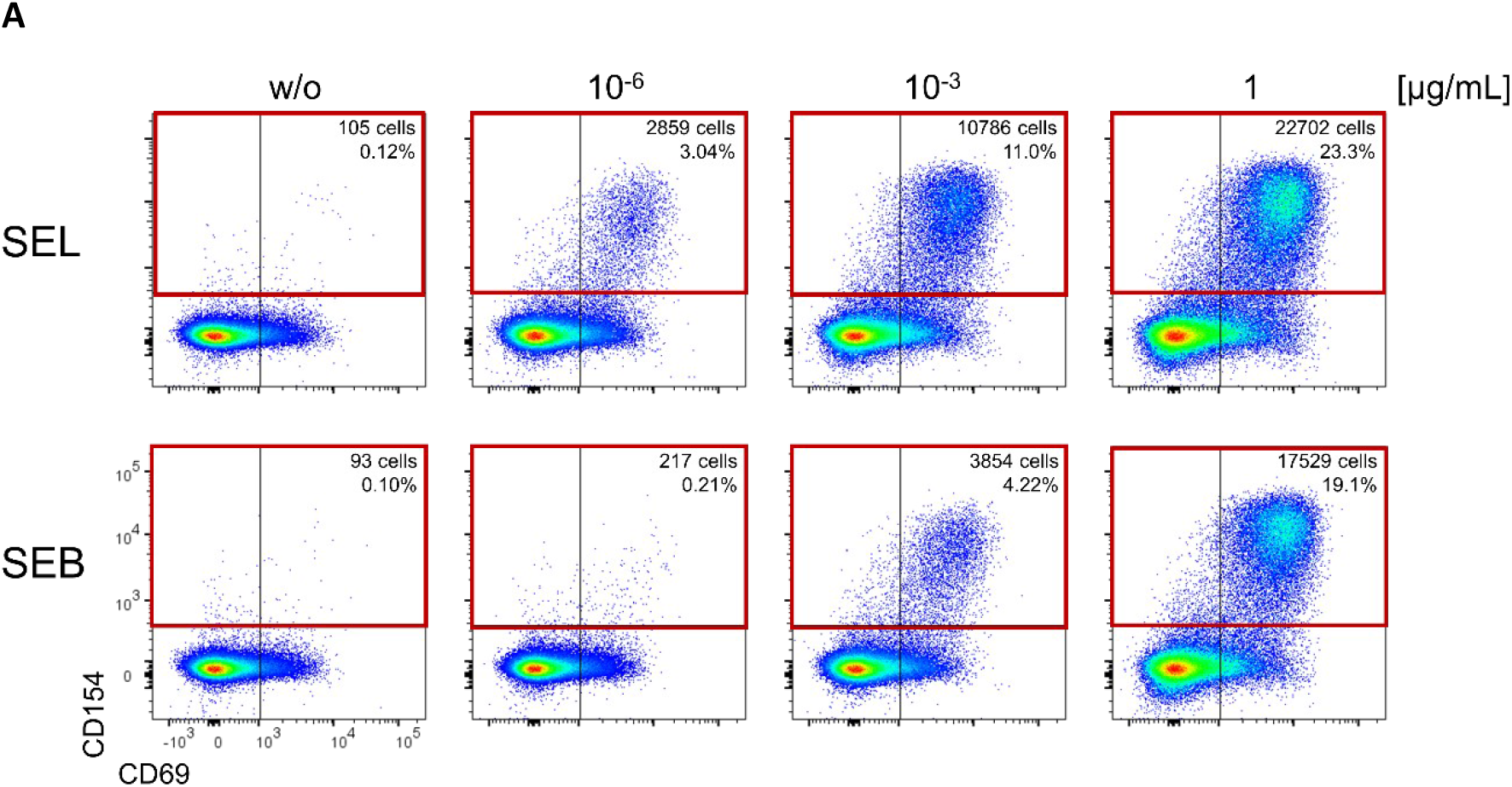
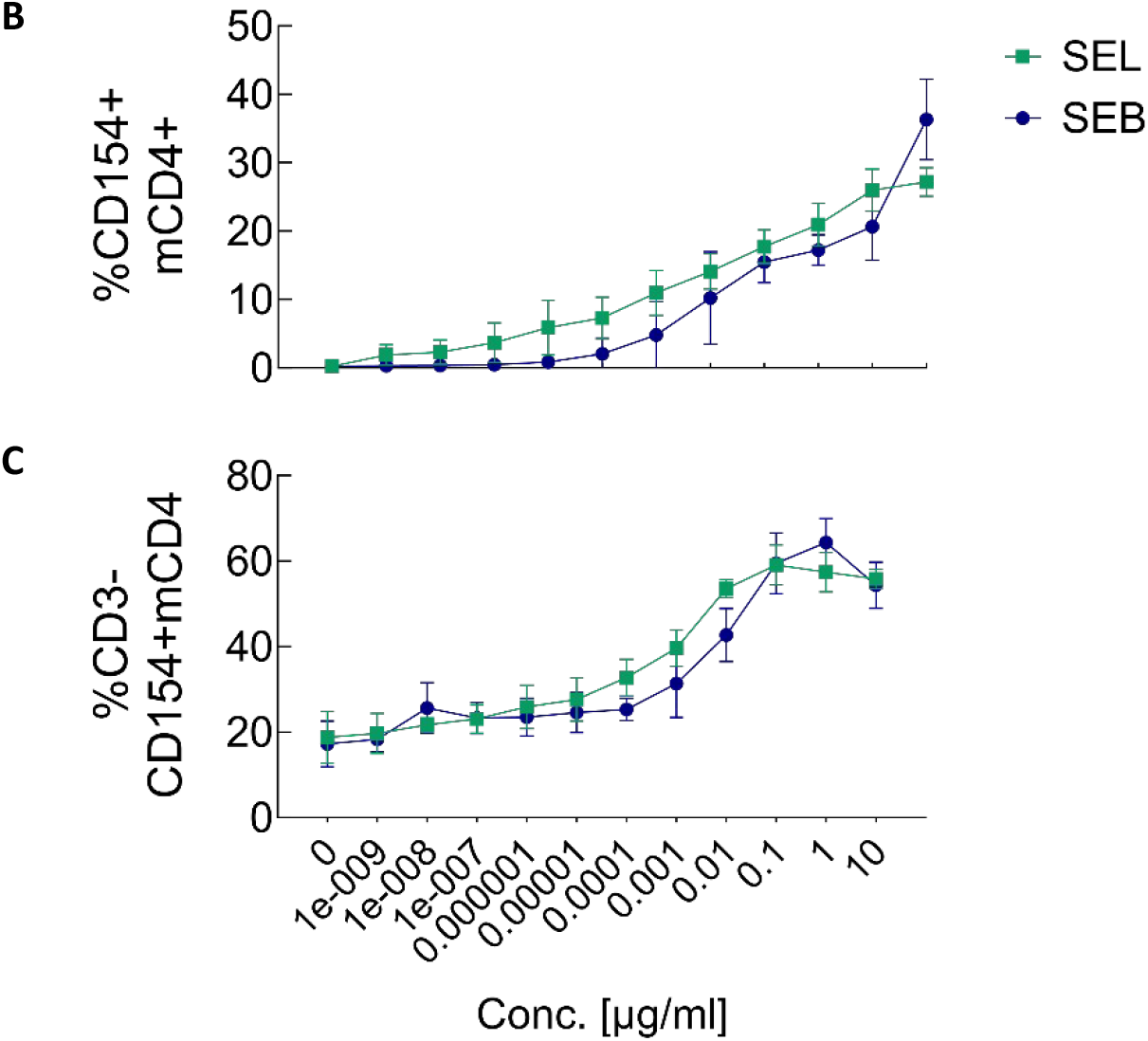
CD4+ T cell activation by SEL and SEB. PBMCs were incubated with the indicated SEL and SEB concentrations for 5 hours and analyzed by flow cytometry for activation marker upregulation. **A**. Example dot plots. **B**. Frequencies of SAg-specific CD154+CD4+ memory T cells. **C**. Percentages of the subpopulation that down-regulates T cell lineage marker CD3 (lower panel). Lines indicate the means +/-SD. N = 4 independent experiments.

We also analysed the proportion of CD154+CD4+ memory T cells that down-regulated the CD3 marker, which indicates T cell receptor (TCR) engagement [14]. A larger share of CD3 downregulation occurred on the surface of T cells activated by SEL than with SEB at various concentrations (Fig 5C). A saturation effect was observed for both superantigens - at ∼0.1 μg/mL for SEL and 1 μg/mL for SEB (Fig 5C). At higher concentrations, the proportion of CD3-T cells decreased, indicating bystander effects, where T cells are activated without a specific TCR engagement, e.g., by cytokine-induced activation marker expression. Both frequencies of CD154+CD4+ and non-bystander CD3-CD154+CD4+ memory T cells were fitted to a non-linear regression model, allowing the extrapolation of SEL and SEB EC_50_ values and the respective confidence intervals (CI) (Fig S4, Table S2). Extrapolated EC50 values of SEL were much lower than those of SEB for activated CD3-CD4+ memory T cells (0.00038 vs. 0.0032 μg/mL, respectively), further confirming that SEL induces strong T cell responses even at very low concentrations.

### Epitope maps

Although some SEs show no obvious relationship to others on the amino acid level, most share substantial sequence similarity with one or more other members of this toxin family. This makes the reliable detection of individual SEs in any matrix extremely challenging and there are currently no commercially available reagents/kits for the specific identification of individual SEs – the importance of developing such reagents for use in, for example, foodchain monitoring, cannot be overestimated. SEL, whose gene has been observed in *S. aureus* strains isolated from both FBO and clinical samples, shares high sequence identities and similarities with four other superantigens: SEI, SEK, SEM and SEQ. In an attempt to identify epitopes on SEL that could be targeted for the development of specific binding modules, the positional residue properties from sequence alignments were color coded and then displayed on the structure of SEL to generate epitope maps, providing a clear visual representation of the positions of unique surface regions. This is illustrated in Figure 6 with the example of SEI. After alignment of the SEL and SEI amino acid sequences (Fig 6A), the residues are colored according to the following scheme: when the identical residue occurs at an equivalent position in both sequences, it is colored red; when the residues at equivalent positions differ but share similar chemical properties (such that their exchange in a real binding interface would not be expected to disrupt the interaction), the residue is colored pink; residues at equivalent positions that have different chemical properties are colored green. This color coding is then displayed on the structure (Fig 6B) to visualize potentially specific epitopes - those whose residue property combinations follow a green > pink > red ratio pattern. After repeating the alignment and color coding for each of the four SEL homologues (Fig S5), the color coded sequences were themselves aligned to generate a “total” epitope map - regions following the designated ratio pattern on this map should be unique to SEL and can be used to distinguish this molecule from all four close homologues (Fig 7).

**Fig 6:**
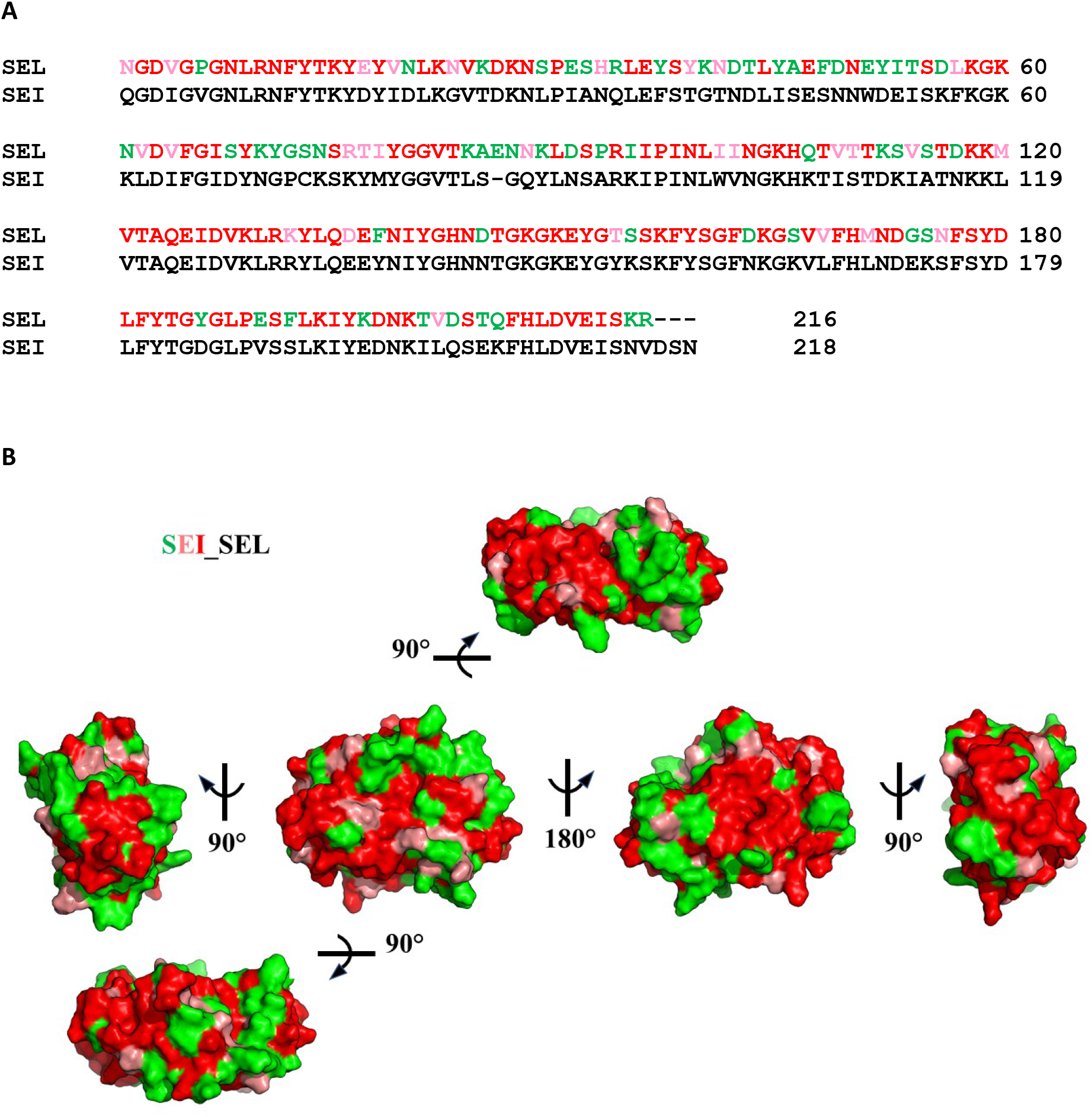
SEL/SEI epitope map. **A**. Alignment of the complete SEL and SEI amino acid sequences with the SEL residues color-coded as described in the text. **B**. Color-coded residues from (**A**) depicted on the model of SEL. Regions comprising residues with a green > pink > red residue profile represent targetable epitopes for differentiating SEL from SEI.

**Fig 7:**
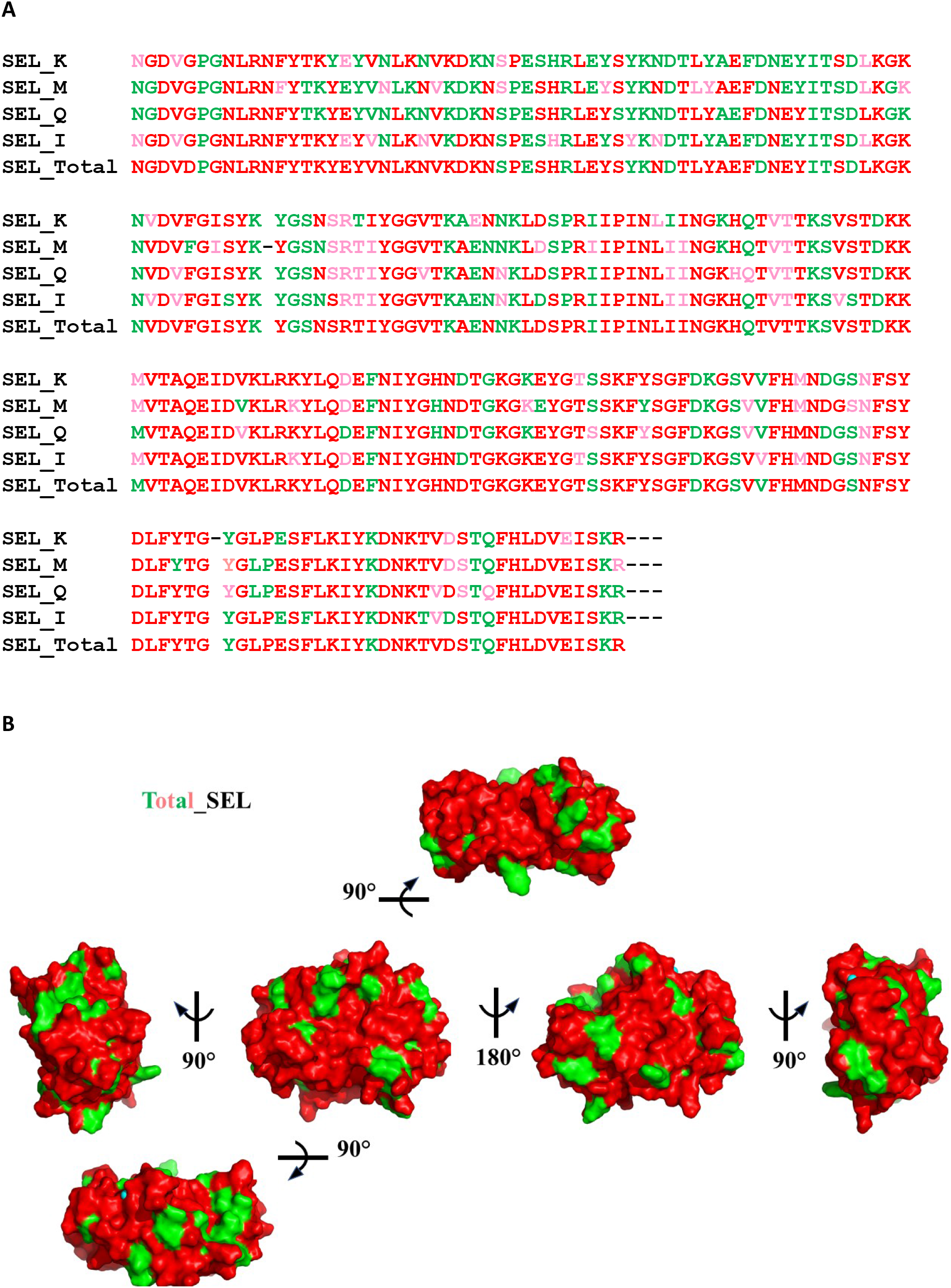
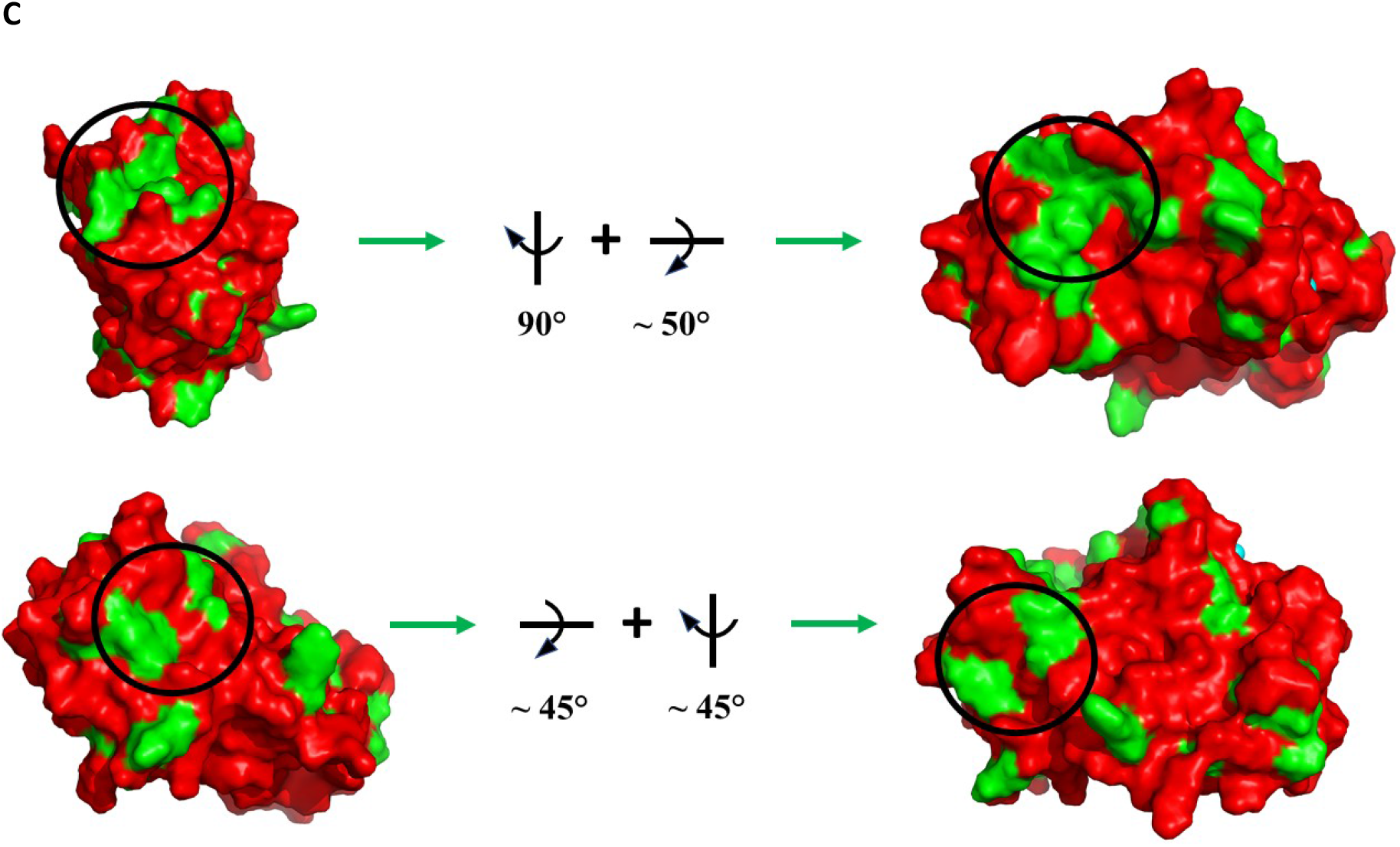
SEL/Group V epitope map. **A**. Sequence alignment of all SEL:Group V SE sequence comparisons, with the consensus color coding (SEL_Total) at the bottom. **B**. Consensus color coding of Group V SEs displayed on the model of SEL, with relative views indicated. The regions indicated in green are those unique for SEL among the Group V SEs. **C**. Multiple views of concave SEL epitopes (beginning with the leftmost view from B) and indicated with black circles. These regions are best suited to binding modules with convex recognition elements, for example nanobodies.

The final three-dimensional epitope map not only shows the areas of specificity but also provides information as to the type of binding module best suited to recognizing each. Because the binding sites of animal-derived antibody molecules comprising heavy and light chains (eg. IgGs) are located in a cleft or groove formed by the CDR loops contributed by both chains, the epitopes that they bind can be crudely described as convex, i.e. they have somewhat protruding surfaces that interact with binding potentials within antibody clefts. Conversely, the binding domains from the heavy chain only antibodies of some groups of animals (cartilaginous fishes and camelids), called nanobodies, have protruding CDR loops that can bind within “concave” epitopes [15]. As indicated on the “total” epitope map shown in Figure 7A and B, in addition to convex epitopes recognizable by standard IgG binders, there are multiple unique epitopes on the surface of SEL with decidedly concave forms that are therefore potential candidates for nanobody binders (Fig 7C). The chemical and steric information from each of these possible epitope regions can, of course, be used to guide their recognition by any type of designed binding module or peptides/aptamers, not only immune system derived molecules.

## Discussion

The potent emetic and immune modulating effects displayed by SEs underscore the urgent need for robust detection assays for their specific identification in foodstuffs and clinical samples – a need complicated by the substantial sequence similarities shared within their subgroups. The ongoing identification of new SE genes, representing a pool of uncharacterized and potentially hazardous agents, adds to this urgency. Further, the occurrence of the gene for SEL in known pathogenicity islands and on mobile genetic elements indicates the potential of its spread via horizontal gene transfer [16]. Indeed, genes for SEL – nearly identical in sequence to that from *S. aureus* - have been detected in isolates of the *Staphylococcal* species *S. argenteus* [17], and *S. epidermis* [18-20] and multiple NCBI entries of genome sequences from *S. pseudintermedius, S. coagulans* and *S. schweizeri* also contain high identity SEL gene sequences. Its not infrequent detection in Staphylococcal isolates and lack of information on its unique features motivated us to undertake a structural evaluation of SEL. The near atomic resolution Xray structure of the toxin reported here provides unambiguous detail of the molecule’s composition, including confirming the presence of a bound Zn ^++^ ion. The high-quality electron density of essentially the complete wildtype amino acid sequence (partly missing only for residue 1 of the chain) also shows the contiguous epitope surface of the molecule that allowed comparison with the closely related Group V SEs, SEK, SEI, SEM and SEQ, for the identification of potential epitopes unique to SEL. The locations of these unique epitopes also inform the choice of binder that can be employed for each, thereby providing the basis information necessary for directed targeting of these regions as a first step in the development of specific diagnostic tests.

The known SE superantigens interact primarily with TCRβ chains, but direct interaction with TCRα chains has also been reported [21]. Our experiments confirmed binding of SEB to TRBV17, but no interaction of SEL with either TRBV17 or TRBV13, in line with earlier findings [7]. Orwin et al. [7] had previously shown that a recombinant SEL gene product from an *S. aureus* isolate obtained from a case of bovine mastitis stimulated proliferation of CD4+ and CD8+ T cells bearing specific TCRβ chains. They additionally demonstrated mitogenicity in a rabbit splenocyte assay at SEL concentrations 100-fold lower than those required to detect similar effects of the toxic shock syndrome toxin (TSST-1). While they did not observe any emetic effect in *M. nemestrina* monkeys, Omoe et al. [6] did observe mild emesis in the related cynomolgus species at the same (100μg/kg) concentration.

To study SEL-induced T cell activation, a CD154-based short-term AIM assay was employed here [11-13]. The advantages of this system include its speed, comprehensiveness and sensitivity. The CD154 marker shows minimal background expression and is upregulated within 5 hours of antigen stimulation, enabling a quantitative readout independent of the variable division speed of different T cell subsets. Compared to the well-characterized superantigen SEB, SEL showed more potent T cell activation, leading to higher frequencies of reactive CD4+ T cells at lower concentrations. These results indicate a higher toxicity of SEL via T cell engagement. In line with earlier observations, CD8+ T cells also participated in the response to SEL [22, 23]. Strong and prolonged T cell responses may lead to the so-called bystander activation, where T cells are activated without a specific TCR engagement, e.g., via cytokines. Since SAgs are capable of activating large fractions of T cells, strong bystander effects may occur at higher concentrations [24, 25]. However, during an antigen-specific T cell response, T cells internalize CD3, a lineage marker and part of the TCR signalling complex [24]. Here, a ratio shift in CD3+ vs CD3-activated CD154+CD4+ memory T cells indicated bystander activation for SEB at the highest SAg concentrations, which may be also supported by binding to other molecular players than TCRs and MHC proteins. For instance, SEB binding to the co-stimulatory molecules CD28 (on T cells) and CD86 or CD40 (on antigen presenting cells) has been described [26-28].

This study reveals unique epitope sites of the Group V Staphylococcal enterotoxin SEL and confirms its extremely potent superantigen activity with human T cells. These results underscore the need and provide the detailed molecular analyses prerequisite for developing precise molecular diagnostic and therapeutic options for identifying and inhibiting this agent.

### Experimental procedures

#### Protein production, crystallization and structure solution

The gene for SEL was amplified via PCR from *S. aureus* strain NCTC10655 provided by the National Reference Laboratory for Coagulase-positive *Staphylococcus aureus* at the German Federal Institute for Risk Assessment, substituting the wildtype N-terminal secretion signal peptide sequence with an affinity tag sequence (amino acid sequence MHHHHHHAGDDDDK) and cloned as an Nco I/Bam HI fragment into vector pET22b. *E. coli* strain BL21(DE3) was transformed with the resulting pSEL construct and protein production induced at an OD_600_ of 0.6 by the addition of 0.1 mM IPTG. The bacteria were harvested 3 hours after induction, pelleted at 5K x g for 10 minutes and the pellets subjected to three freeze/thaw cycles as per Johnson and Hecht [29]. The extracted cell mass was centrifuged at 21K x g for 1 hour, the supernatant treated with Benzonase for 1 hour at RT followed by filtration (0.45μm), and then subjected to Ni-NTA affinity chromatography followed by size exclusion on a Superdex S75 (GE Healthcare) prep column. The resulting peak fractions were analyzed via SDS-PAGE, appropriately pooled and concentrated (Amicon) and subjected to crystallization screening.

Fine screens were prepared based on initial hits from commercial screens (Jena Biosciences) and diffracting crystals obtained in hanging drops at 20°C in a mother liquor consisting of 12% PEG 8000, 100 mM imidazole, pH 8.0. Drops consisted of a 2:1 ratio of protein solution:mother liquor. Crystals reproducibly appeared as clusters of large (ca. 0.4 mm) and extremely mechanically robust plates. Diffraction screening was performed on Beamline 14.3 of the BESSY Synchrotron at the Helmholtz Zentrum Berlin at RT using the HC-1 humidity stream [30] and diffracting crystals flash frozen without cryoprotectant via automated switching from the humidity stream to a cryostream. Complete diffraction data sets were then collected also at Beamline 14.3. The data were processed using xdsapp [31] and the structure was solved by molecular replacement using 3EA6 (SEK) as the search model with the program Phaser [32] followed by model building and refinement with COOT [33] and Phenix [34]. All images were generated using PyMOL (The PyMOL Molecular Graphics System, Version 1.2r3pre, Schrödinger, LLC).

#### T cell activation studies

Human buffy coats containing T cells and written permission to use them for the experiments described, in accordance with the declaration of Helsinki principles, were obtained from the German Red Cross (Votum EA1_217_19, ethics committee of the Charité – Universitätsmedizin Berlin).

PBMCs were isolated by standard Ficoll-Paque plus TM (Cytiva) density gradient centrifugation and cultured in complete RPMI 1640 (PAN Biotech)-based T cell medium supplemented with 5% heat-inactivated human serum AB (PAN Biotech) in flat-bottom plates at 2.5.10^6^ cells/cm^2^. Cells were either left unstimulated (w/o) or incubated with the indicated concentrations of SEL (previously described) or SEB (Sigma-Aldrich) in the presence of CD40 blocking antibody (Biolegend, clone HB14) [11, 12]. After 5 h, cells were stained for surface markers and analysed on a BD FACSAria III sorter (BD Biosciences) equipped with three lasers and 14 fluorescence detectors. Flow cytometry data were further analysed using FlowJo Software (Version 10.10.0) and graphs were designed and analysed in GraphPad Prism (Version 10.1.2).

## Data availability

The SEL structural data have been deposited in the Protein Databank (www.rcsb.org) under PDB ID 9Q9U.

## Supporting information

This article contains supporting information.

## Acknowledgments

Diffraction data was collected at beamline 14.3 at the BESSY II electron storage ring operated by the Helmholtz-Zentrum Berlin für Materialien und Energie; we gratefully acknowledge Manfred Weiss and Frank Lennartz for support at the beamline and in particular Laila Benz and Gert Weber for assistance with the HC-1 dehydration device. We additionally thank Josephine Becker for technical assistance and acknowledge Jens-André Hammerl for assistance during the freezing of crystals.

## Author contributions

SFM initiated and coordinated the study, cloned, produced and purified the toxin, performed initial FACS analyses confirming T cell activation, crystallized the protein, measured diffraction data, solved and refined the structure, performed all structural and epitope analyses, prepared figures and wrote the manuscript.

KS supervised the T cell activation studies, analysed FACS data, prepared figures and wrote the manuscript.

CC performed the T cell activation studies, analysed FACS data, prepared figures and wrote the manuscript.

EK produced and purified toxin, prepared materials for initial FACS analyses and wrote the manuscript.

## Funding and additional information

This work was supported via grants from the German Federal Ministry for Food and Agriculture: BfR-BIOS-23-1322-839 to SFM and BfR-CPS-23-1322-848 to CC and KS.

## Abbreviations

The abbreviations used are:

*S. aureus*: (*Staphylococcus aureus*)
spp.: (species)
SE: (*Staphylococcal* enterotoxin)
SEL: (*Staphylococcal* enterotoxin L)
(SAg): superantigen,
TCRα: (T cell receptor alpha chain)
TCRβ: (T cell receptor beta chain)
(AIM) assay: activation-induced marker

