## Supplemental Figures and Tables for "Identification of targetable epitope surfaces from the high resolution structure of the superantigen *Staphylococcal* Enterotoxin L"

File contents:

Supplementary Figures:

S1: **The structure of SEL at 1.3 Å**
S2: **Gating strategy**
S3: **Expression of further activation markers by CD4+ and CD8+ T Cells in the AIM assay**
S4: **Non-linear regression curves of SEL and SEB antigen-specific T cell responses**
S5: **Group V SE sequence comparisons with SEL**

Supplementary Tables:

Table S1: **Data collection and refinement statistics (molecular replacement)**
Table S2: **Non-linear regression model of SEL and SEB-induced T cell responses**

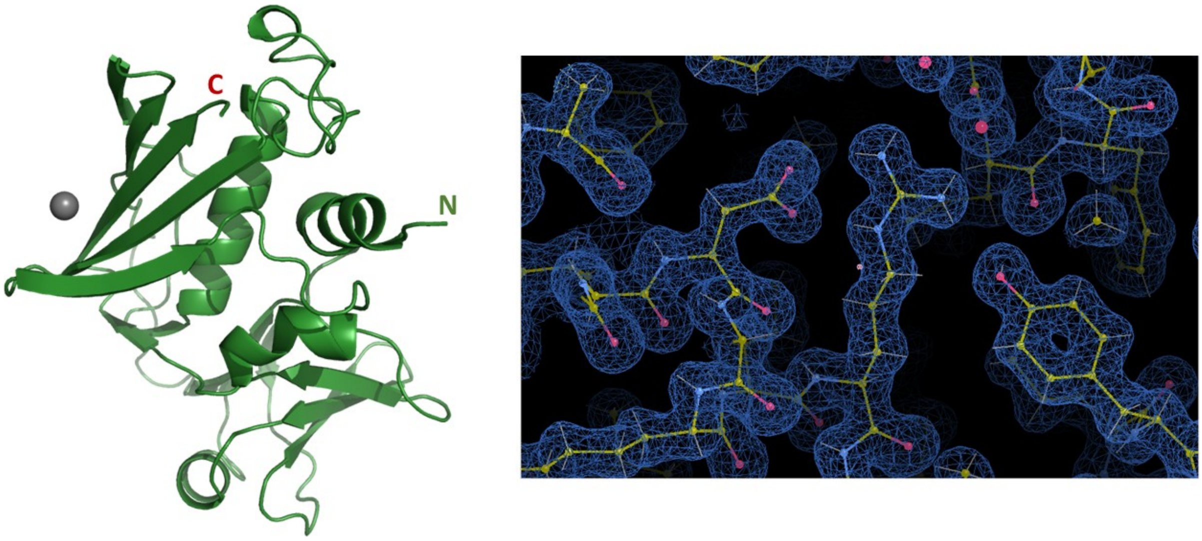

Fig S1. **The structure of SEL at 1.3 Å.** The left panel shows SEL rendered as ribbons with the N and C termini explicitly marked; the TCR binding site is on the right side of the molecule, the MHC II binding site is on the left side, at the position where a bound $Zn^{++}$ atom is depicted as a grey sphere. The right panel shows representative electron density contoured at 1.0 $\delta$.

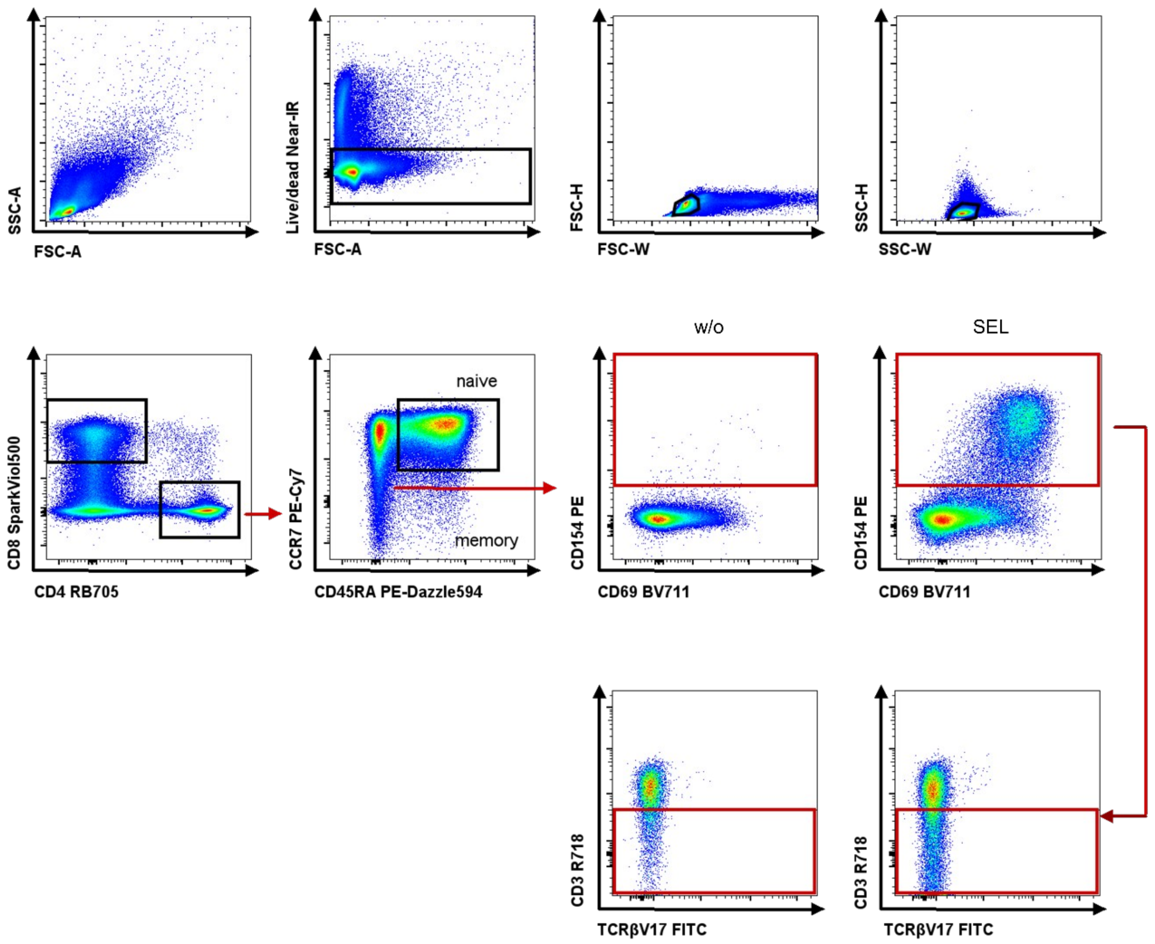

Fig S2: **Gating strategy.** PBMCs were stained with up to 13 fluorochrome-labelled antibodies and a dead cell stain. Sequential gating was performed on live and single cells, followed by gating on CD4+ or CD8+ memory T cells (non-naïve, naïve defined as CD45RA+CCR7+). Expression of activation markers including CD154, CD69 or CD137 was assessed. For CD154+ cells, further analysis assessed down-regulation of CD3 and the presence of Vb 17 (TRVB19) among activated T cells. Dot plots from a representative experiment with SEL [1 µg/mL].

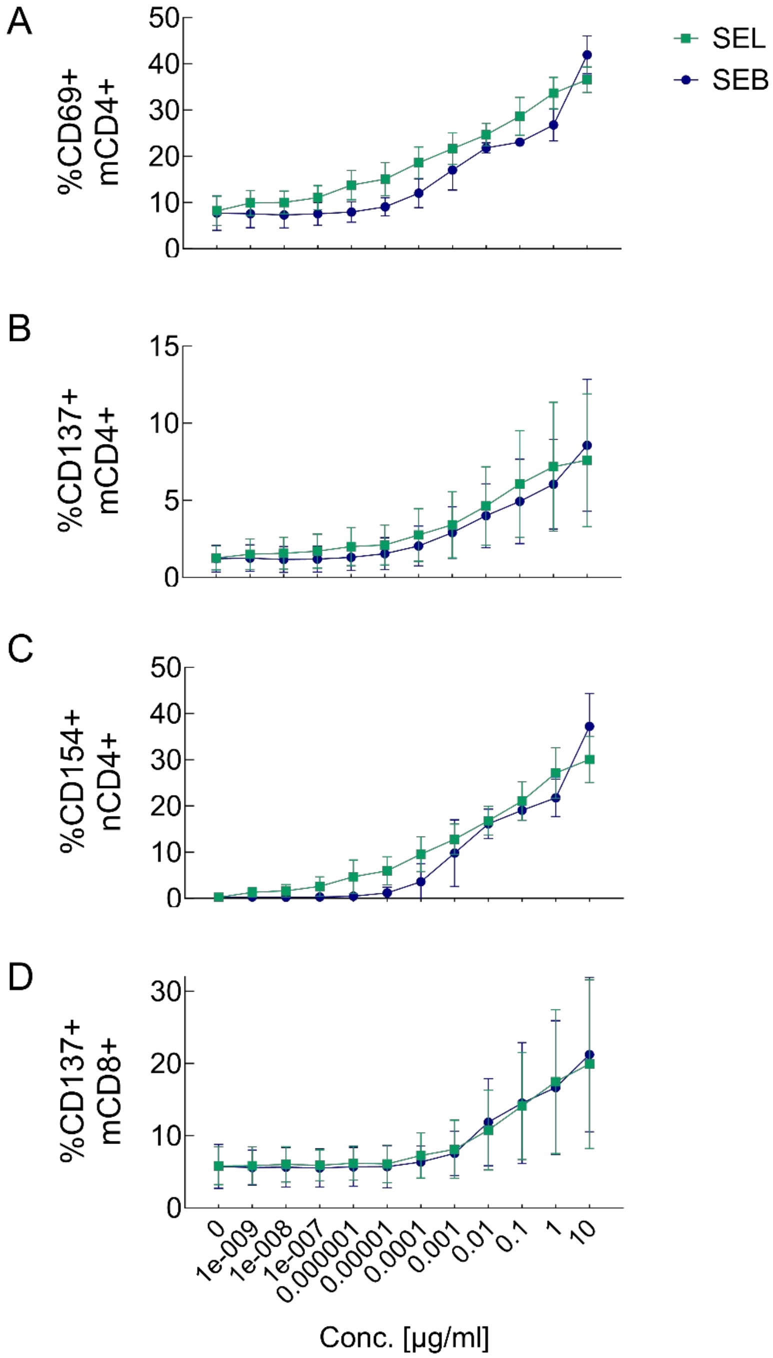

Fig S3: **Expression of further activation markers by CD4+ and CD8+ T Cells in the AIM assay.** PBMCs were incubated with the indicated SEL and SEB concentrations for 5 hours and analyzed by flow cytometry for upregulation of surface markers. **(A)** CD69 and **(B)** CD137 on CD4+ memory T cells, **(C)** CD154 on CD4+ naive T cells and **(D)** CD137 on CD8+ memory T cells. Lines indicate the means +/-SD. N = 4 independent experiments.

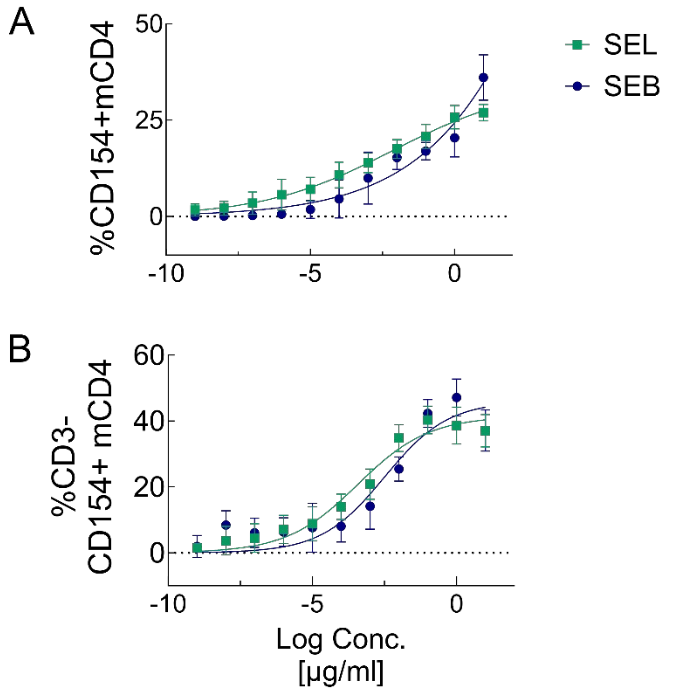

Fig S4: **Non-linear regression curves of SEL and SEB antigen-specific T cell responses.** Depicted are non-linear regression curves for the extrapolation of EC50 values and confidence intervals of SEL- and SEB-specific T cell responses (see also **Table S2**). Frequencies of **(A)** CD154+mCD4+ T cells and **(B)** the subpopulation that down-regulated the T cell lineage marker CD3 were used to fit the regression model after subtraction of background T cell activation without antigen stimulation. N = 4 independent experiments.

```
SEL      NGDVGPGNLRNFYTK YEYVNLKNVKDKNSPESHRLEYSYKNDTLYAEFDNEYITSDLKGK   60
SEK      QGDIGIDNLRNFYTKKDFVDLKDVKDNDTPIANQLQFSNESYDL ISESKDFNKFSNFKGK   60

SEL      NVDVFGISYKYGSNSRTIYGGVTKAENNKLDSPRIIPINLIINGKHQTVTTKSVSTDKKM   120
SEK      KLDVFGISYNGQCNTKYIYGGVTA-TNEYLDKSRNIPINIWINGNHKTISTNKVSTNKKF   119

SEL      VTAQEIDVKLRKYLQDEFNIYGHNDTGKGKEYGTSSKFYSGFDKGSVVFHMNDGSNFSYD   180
SEK      VTAQEIDVKLRKYLQEEYNIYGHNGTKKGEEYGHKSKFYSGFNIGKVTFHLNNNDTFSYD   179

SEL      LFYTG-YGLPESFLKIYKDNKTVDSTQFHLDVEISKR---      216
SEK      LFYTGDDGLPKSFLKIYEDNKTVESEKFHLDVDISYKETI      219

>>>

SEL      NGDVGPGNLRNFYTKYEYVNLKNVKDKNSPESHRLEYSYKNDTLYAEFDNEYITSDLKGK 60
SEM      --DVGVLNLRNYYGSYPIESHQNINPDNNSLSHQLVFSKDNSTVTAEFKNVEDVKKFKNR 58

SEL      NVDVFGISYK-YGSNSRTIYGGVTKAENNKLDSPRIIPINLIINGKHQTVTTKSVSTDKK 119
SEM      AVDVYGLSYSGYCLKNKYMYGGVTLA-GDYLEKSRCIPINLWVNGNHKTISTDKVSTNKK 117

SEL      MVTAQEIDVKLRKYLQDEFNIYGHNDTGKGKEYGTSSKFYSGFDKGSVVFHMNDGSNFSY 179
SEM      IVTAQEIDTKLRRYLQEEYNIYGFNDTNKGRNYGTKSKFFSGFNTGKISFHLNDGTSFSY 177

SEL      DLFYTGYGLPESFLKIYKDNKTVDSTQFHLDVEISKR---      216
SEM      DLFDTGTGQAESFLKIYNDNKTVETDKFHLDVEISYKDES      217

>>>

SEL      NGDVGPGNLRNFYTKYEYVNLKNVKDKNSPESHRLEYSYKNDTLYAEFDNEYITSDLKGK 60
SEQ      --DVGVINLRNFYANYEPEKLQGVSSGNFSTSHQLEYIDGKYTLYSQFHNEYEAKRLKDH 58

SEL      NVDVFGISYKYGSNSRTIYGGVTKAENNKLDSPRIIPINLIINGKHQTVTTKSVSTDKKM 120
SEQ      KVDIFGISYSGLCNTKYMYGGITLA-NQNLDKPRNIPINLWVNGKQNTISTDKVSTQKKE 117

SEL      VTAQEIDVKLRKYLQDEFNIYGHNDTGKGKEYGTSSKFYSGFDKGSVVFHMNDGSNFSYD 180
SEQ      VTAQEIDIKLRKYLQNEYNIYGFNKTKKGQEYGYQSKFNSGFNKGKITFHLNNEPSFTYD 177

SEL      LFYTGYGLPESFLKIYKDNKTVDSTQFHLDVEISKR---       216
SEQ      LFYTGTGQAESFLKIYDDNKTIDTENFHLDVEISYEKTE       216

>>>

SEL      NGDVGPGNLRNFYTKYEYVNLKNVKDKNSPESHRLEYSYKNDTLYAEFDNEYITSDLKGK 60
SEI      QGDIGVGNLRNFYTKYDYIDLKGVTDKNLPIANQLEFSTGTNDLISESNNWDEISKFKGK 60

SEL      NVDVFGISYKYGSNSRTIYGGVTKAENNKLDSPRIIPINLIINGKHQTVTTKSVSTDKKM 120
SEI      KLDIFGIDYNGPCKSKYMYGGVTLS-GQYLNSARKIPINLWVNGKHKTISTDKIATNKKL 119
```

```
SEL      VTAQEIDVKLRKYLQDEFNIYGHNDTGKGKEYGTSSKFYSGFDKGSVVFHMNDGSNFSYD 180
SEI      VTAQEIDVKLRRYLQEEYNIYGHNNTGKGKEYGYKSKFYSGFNKGKVLFHLNDEKSFSYD 179

SEL      LFYTGYGLPESFLKIYKDNKTVDSTQFHLDVEISKR---       216
SEI      LFYTGDGLPVSSLKIYEDNKILQSEKFHLDVEISNVDSN       218

>>>

SEL_K     NGDVGPGNLRNFYTKYEYVNLKNVKDKNSPESHRLEYSYKNDTLYAEFDNEYITSDLKGK 60
SEL_M     NGDVGPGNLRNFYTKYEYVNLKNVKDKNSPESHRLEYSYKNDTLYAEFDNEYITSDLKGK
SEL_Q     NGDVGPGNLRNFYTKYEYVNLKNVKDKNSPESHRLEYSYKNDTLYAEFDNEYITSDLKGK
SEL_I     NGDVGPGNLRNFYTKYEYVNLKNVKDKNSPESHRLEYSYKNDTLYAEFDNEYITSDLKGK
SEL_Total NGDVDPGNLRNFYTKYEYVNLKNVKDKNSPESHRLEYSYKNDTLYAEFDNEYITSDLKGK

SEL_K     NVDVFGISYK YGSNSRTIYGGVTKAENNKLDSPRIIPINLIINGKHQTVTTKSVSTDKK
SEL_M     NVDVFGISYK-YGSNSRTIYGGVTKAENNKLDSPRIIPINLIINGKHQTVTTKSVSTDKK
SEL_Q     NVDVFGISYK YGSNSRTIYGGVTKAENNKLDSPRIIPINLIINGKHQTVTTKSVSTDKK
SEL_I     NVDVFGISYK YGSNSRTIYGGVTKAENNKLDSPRIIPINLIINGKHQTVTTKSVSTDKK
SEL_Total NVDVFGISYK YGSNSRTIYGGVTKAENNKLDSPRIIPINLIINGKHQTVTTKSVSTDKK

SEL_K     MVTAQEIDVKLRKYLQDEFNIYGHNDTGKGKEYGTSSKFYSGFDKGSVVFHMNDGSNFSY
SEL_M     MVTAQEIDVKLRKYLQDEFNIYGHNDTGKGKEYGTSSKFYSGFDKGSVVFHMNDGSNFSY
SEL_Q     MVTAQEIDVKLRKYLQDEFNIYGHNDTGKGKEYGTSSKFYSGFDKGSVVFHMNDGSNFSY
SEL_I     MVTAQEIDVKLRKYLQDEFNIYGHNDTGKGKEYGTSSKFYSGFDKGSVVFHMNDGSNFSY
SEL_Total MVTAQEIDVKLRKYLQDEFNIYGHNDTGKGKEYGTSSKFYSGFDKGSVVFHMNDGSNFSY

SEL_K     DLFYTG-YGLPESFLKIYKDNKTVDSTQFHLDVEISKR---
SEL_M     DLFYTG YGLPESFLKIYKDNKTVDSTQFHLDVEISKR---
SEL_Q     DLFYTG YGLPESFLKIYKDNKTVDSTQFHLDVEISKR---
SEL_I     DLFYTG YGLPESFLKIYKDNKTVDSTQFHLDVEISKR---
SEL_Total DLFYTG YGLPESFLKIYKDNKTVDSTQFHLDVEISKR
```

Fig S5: **Group V SE sequence comparisons with SEL.** Individual comparisons of each Group V SE sequence with that of SEL. The comparison sequences are in black with the corresponding color coded residues of the SEL sequence shown above them. The “Total” or consensus color coded sequence is shown last.

**Table S1 Data collection and refinement statistics (molecular replacement)**

| | SEL |
|---|---|
| **Data collection** | |
| Space group | P 1 21 1 |
| Cell dimensions | |
| *a*, *b*, *c* (Å) | 32.3 , 63.86 , 48.62 |
| α, β, γ (°) | 90, 92.33, 90 |
| Resolution (Å) | 1.31 (1.31 – 1.39) * |
| $CC_{1/2}$ | 0.993 (0.228) |
| $I / \sigma I$ | 6.61 (1.63) |
| Completeness (%) | 87.48 (47.01) |
| Redundancy | 3.23 (2.68) |
| **Refinement** | |
| Resolution (Å) | 1.31 - 28.8 |
| No. reflections | 41477 (2772) |
| $R_{\text{work}}$ / $R_{\text{free}}$ | 0.1233 / 0.1596 |
| No. atoms | |
| Protein | 1739 |
| Ligand/ion | 1 |
| Water | 153 |
| *B*-factors (Å$^2$) | |
| Protein | 21.90 |
| Ligand/ion | 13.99 |
| Water | 37.60 |
| R.m.s. deviations | |
| Bond lengths (Å) | 0.007 |
| Bond angles (°) | 0.94 |

*The complete dataset was measured from a single crystal; values in parentheses are for highest-resolution shell.

Table S2. **Non-linear regression model of SEL and SEB-induced T cell responses**. Frequencies of CD154+CD4+ or CD3-CD154+CD4+ memory T cells after subtraction of background T cell activation without antigen stimulation (w/o) were fitted in a non-linear regression model ($R^2$>0.90). The table summarizes EC50 values and confidence intervals (CI). N = 4 independent experiments.

| log(agonist) vs. response -- Variable slope (four parameters) | **CD154+CD4+** (background corrected) | | **CD3-CD154+CD4+** (background corrected) | |
|---|---|---|---|---|
| | **SEL** | **SEB** | **SEL** | **SEB** |
| **Best-fit values** | | | | |
| Bottom | 0 | 0 | 0 | 0 |
| Top | 34.16 | 228.1 | 41.47 | 46.01 |
| LogEC50 | -2.154 | 5.234 | -3.426 | -2.495 |
| HillSlope | 0.1932 | 0.1763 | 0.3519 | 0.3947 |
| **EC50** | **0.007013** | ---[¥] | **0.0003754** | **0.003197** |
| Span | 34.16 | 228.1 | 41.47 | 46.01 |
| **95% CI** (profile likelihood) | | | | |
| Top | 30.70 to 39.51 | ---[¥] | 36.11 to 50.75 | 35.23 to 271.2 |
| LogEC50 | -2.736 to -1.325 | -1.220 to ---[¥] | -4.065 to -2.514 | -3.623 to 6.746 |
| HillSlope | 0.1675 to 0.2212 | 0.1300 to 0.3239 | 0.2169 to 0.5929 | 0.1124 to 1.325 |
| EC50 [µg/ml] | 0.001836 to 0.04727 | 0.06028 to ---[¥] | 8.612e-005 to 0.003064 | 0.0002382 to 5575613 |
| **Goodness of Fit** | | | | |
| Degrees of Freedom | 8 | 8 | 8 | 8 |
| **R squared** | **0.9973** | **0.9657** | **0.97** | **0.9033** |
| Sum of Squares | 2.337 | 45.86 | 73.21 | 261.6 |
| Sy.x | 0.5405 | 2.394 | 3.025 | 5.719 |
| Constraints | | | | |
| Bottom | Bottom = 0 | Bottom = 0 | Bottom = 0 | Bottom = 0 |
| Number of points | | | | |
| # of X values | 11 | 11 | 11 | 11 |
| # Y values analyzed | 11 | 11 | 11 | 11 |

[¥] The curve of CD154+CD4+ memory T cells upon SEB stimulation did not reach saturation and therefore EC50 and CI could not be calculated by the regression model.
